# Annotated Compendium of 102 Breast Cancer Gene-Expression Datasets

**DOI:** 10.1101/2023.09.22.559045

**Authors:** Ifeanyichukwu O. Nwosu, Daniel D. Tabler, Greg Chipman, Stephen R. Piccolo

## Abstract

Transcriptomic data from breast-cancer patients are widely available in public repositories. However, before a researcher can perform statistical inferences or make biological interpretations from such data, they must find relevant datasets, download the data, and perform quality checks. In many cases, it is also useful to normalize and standardize the data for consistency and to use updated genome annotations. Additionally, researchers need to parse and interpret metadata: clinical and demographic characteristics of patients. Each of these steps requires computational and/or biomedical expertise, thus imposing a barrier to reuse for many researchers. We have identified and curated 102 publicly available, breast-cancer datasets representing 17,151 patients. We created a reproducible, computational pipeline to download the data, perform quality checks, renormalize the raw gene-expression measurements (when available), assign gene identifiers from multiple databases, and annotate the metadata against the National Cancer Institute Thesaurus, thus making it easier to infer semantic meaning and compare insights across datasets. We have made the curated data and pipeline freely available for other researchers to use. Having these resources in one place promises to accelerate breast-cancer research, enabling researchers to address diverse types of questions, using data from a variety of patient populations and study contexts.

## Background

Among women worldwide, breast cancer is the most frequently diagnosed cancer type and the leading cause of cancer death; it accounts for 23% of total cases and 14% of deaths^1^, with over 2 million new cases diagnosed and 600,000 deaths in 2020^2^. This is a huge public health burden. Breast cancers exhibit considerable diversity and complexity^3^ and are driven by a combination of factors, including genomic material inherited from parents and an accumulation of somatic events within individual cells; these events include mutations, chromosomal rearrangements, gene amplifications, and deletions that eventually lead to evasion of apoptosis and unregulated cellular proliferation^4^. While earlier diagnoses and treatment advances have improved survival and lowered mortality rates^5,6^, researchers continue to seek to understand the molecular underpinnings of this disease, identify subtypes of the disease, and explore ways to tailor treatments for individual patients^3,7^.

Microarrays and RNA sequencing have been used in hundreds of thousands of gene-expression studies (spanning many biomedical contexts)^8^, helping to identify differentially expressed genes, define disease subtypes, inform treatment and patient-management strategies, predict patient outcomes, etc.^9^ Much of the data from these studies has been deposited in public databases such as Gene Expression Omnibus (GEO)^8,10^ and ArrayExpress^11^. These databases offer opportunities for reuse, including studies that combine insights from multiple datasets. However, making data accessible is not enough. To be most useful for addressing biomedical questions, the data must also be findable, interoperable, and reusable^12^. First, researchers must locate the data. Gene-expression datasets are available in diverse repositories. These repositories support keyword searches; however, our experience is that these searches miss many relevant datasets and require considerable manual effort to filter through search results. Although in some cases researchers can reuse data exactly in the form they are found in public repositories, data depositors use a heterogeneous mix of methods for verifying data quality, normalizing expression measurements, and then filtering, summarizing, and annotating these measurements. Sometimes steps are skipped or may be completed using invalid methods. Furthermore, genome annotations are regularly updated, so datasets that were annotated years ago may have stale annotations. Fortunately, gene-expression data are frequently provided in raw form, so researchers can perform these steps on their own, thus resulting in datasets that are more consistent and of higher quality.

A key challenge to reusing publicly available datasets is working with metadata. Nearly all gene-expression data from human patients have clinical and/or demographic data associated with them. However, researchers use different terms, abbreviations, and spellings (including misspellings) to describe the same clinical and demographic phenomena. For example, we have seen variable names as diverse as "drfs_even_time_years", "distal_recurrence_time", "distant_recurrence_free_survival_mo", "dmfs_time", "t_tdm", and "time" to describe the length of time a patient survived without a metastatic event after diagnosis. Some variable names are short and/or vague (e.g., "cell_type", "chemo", "nhg", "pt"). Sometimes measurement units are included in names of numeric variables (e.g., "age_years", "size_mm", "mfsdel_month"), but other times they are not. Some variable names incorrectly or inadequately reflect the semantics of those variables. For example, "ethnicity" sometimes refers to population ancestral groups (often referred to as "race"); "gender" sometimes refers to a person’s biological sex assigned at birth. Clarifying these semantic inconsistencies is especially important when researchers wish to identify datasets that have particular metadata variables and/or to combine multiple datasets.

Formatting differences also cause challenges. In GEO, data are provided in the "series matrix" and SOFT formats^13^, whereas ArrayExpress uses standardized formats like MAGE-TAB^14^. Software tools are available to download and convert the data into formats and data structures that are not specific to gene-expression data^15,16^; however, other challenges remain. Researchers sometimes store multiple values in a single cell (e.g., "female|tamoxifen|58"), use key/value pairs within cells (e.g., "sex: female", "treatment=tamoxifen", "age=58 years"), or include missing values (using descriptors such as "NA", "n/a", or "?")^17^. Addressing these problems requires time and expertise. To ensure semantic consistency, biomedical domain experts must manually review, interpret, and standardize metadata. To deal with formatting problems, researchers often must write custom computer code. These barriers have caused these valuable data resources to be underutilized and have slowed the research enterprise.

We have addressed these challenges for breast cancer by finding and curating 102 gene-expression datasets from the public domain. These datasets represent 17,151 patient samples. For each dataset, we performed quality-control checks, uniformly processed the expression data, mapped the expression data to multiple gene-reference databases, and standardized the metadata variables against the National Cancer Institute Thesaurus (NCIT)^18^, a popular reference standard with unique codes and synonyms for biomedical terms. Additionally, a practicing medical oncologist reviewed ambiguous term mappings. We have deposited these datasets, along with the code we used to process them, in an open-access repository, thus making it easy for other researchers to find, access, integrate, and reuse the data, as well as to reproduce our process. This repository is not tied to any programming language—the data can be imported using a variety of tools, including Microsoft Excel and software libraries available for commonly used programming languages.

## Methods

### Data finding

Initially, we searched for datasets within GEO and ArrayExpress. Our initial research interest was the interaction between race and HER2 amplification status among breast-cancer patients, and this guided our choice of search parameters. For GEO, our search parameters were ("Breast Cancer"[All Fields] AND "Her2"[All Fields] AND "race"[All Fields]). We selected the study types as “Expression profiling by array” or “Expression profiling by high-throughput sequencing”. We individually screened the available datasets for relevance and duplicates. In ArrayExpress, our search parameters were "Breast Cancer" AND "Her2" AND "race". Next, we searched within PubMed for articles that matched the following criteria: "Breast Cancer" AND "Her2" AND "race". These criteria resulted in a limited number of datasets, so we expanded our searches in PubMed to breast cancer more broadly. We also searched for datasets using the Google search engine. As part of this process, we identified two existing collections that included breast-tumor expression data: MetaGxData^19^ and curatedBreastData^20^. We used these collections as resources for identifying relevant breast-cancer datasets. Finally, we examined cancer-genomics repositories for additional datasets. The International Cancer Genome Consortium (ICGC) data portal^21,22^ contained five breast-cancer datasets, two of which provided gene-expression data; raw data were available for one of them. cBioPortal for Cancer Genomics^23^ provided a copy of gene-expression data from the Molecular Taxonomy of Breast Cancer International Consortium (METABRIC) project^24^. The Cancer Genome Atlas (TCGA) provided gene-expression data for many cancer types, including breast cancer^25^. We used a reprocessed version of the data stored in GEO^26^.

We found a mix of different data-generation platforms, including Affymetrix, Agilent and Illumina microarrays, custom microarrays, as well as Illumina-based RNA sequencing. Because the Affymetrix platforms were the most common—and well-recognized normalization and annotation packages could be applied to these platforms—we focused on these. We also included datasets that had been processed using RNA sequencing technology because of its advantages over microarray technologies^27^. With one exception, we excluded datasets that used Agilent microarrays, Illumina microarrays, or uncommon microarray platforms to reduce heterogeneity. The METABRIC project used Illumina microarrays; we included this dataset in our compendium because it is large (1980 samples) and provides extensive metadata. We excluded Affymetrix datasets for which raw data were unavailable or that were from cell lines or single cells. Raw data were not publicly available for the METABRIC dataset or the RNA-sequencing datasets; we used the processed versions of these datasets. This resulted in 102 datasets with a total of 17,151 samples. Most of the samples are from breast tumors; however, a small number are from normal breast tissue as indicated in the metadata.

### Data curation

We automated all steps of the curation process using custom computer scripts. Some scripts used the bash interpreter (Linux operating system); other scripts used the R statistical software^28^. We configured the scripts to be executed within Docker containers so that our curation process is reproducible and portable across computer systems^29^. We obtained the GEO data using the GEOquery^15^ package. For the other sources, our scripts downloaded the data directly from the repositories. The data comprised both gene-expression data and metadata, which we processed to achieve consistency across all datasets.

Our workflow was divided into six major steps described below.

In step one, we downloaded the metadata for Affymetrix datasets, cleaned up column names by removing non-informative suffixes (e.g "_ch1") and removing columns that we determined would not add value in downstream analyses. For example, some columns had only one unique value (e.g., "extraction protocol", "last_update_date"), while others had all unique values such (e.g., "sample_name"). Because this process was subjective, and other researchers might find these columns useful, we retained the unfiltered metadata as a separate part of the outputs. Upon reviewing the metadata, we made additional customizations on a case-by-case basis. When a single dataset was processed using more than one platform, we separated them into multiple data files, according to their respective platforms. Finally, we summarized the metadata variables from each dataset. These summaries indicated the unique values for each variable, the number of unique values, and the minimum, maximum and mean (for numeric variables).

In step two, we downloaded raw CEL files for the Affymetrix datasets and normalized them using the SCAN.UPC^30^ package, yielding zero-centered, log2-transformed intensities. This normalization step made use of custom chip definition files (CDF) for Affymetrix GeneChips using updated probe-set definitions from the Brainarray Microarray consortium^31^. Custom CDFs are necessary because the manufacturer’s probe-set definitions have become outdated and are sometimes incorrectly assigned to genes or to multiple genes. Some GEO datasets had a mixture of Affymetrix and Agilent data. Our scripts selected the Affymetrix samples only.

In step three, we performed quality-control measures on the expression data. We used the DoppelgangR^32^ package to identify Affymetrix samples that may have been duplicated. Duplicates can occur when samples are inadvertently mislabeled or when studies include samples from previously published datasets. After manually reviewing the sample pairs that had been flagged as duplicates by *DoppelgangR*, we determined that an expression-similarity threshold of 0.99 may be appropriate for excluding samples; this score errs on the side of only excluding samples that were most likely to be duplicates. The sample pairs that exceed this threshold are indicated in our output files. However, we retained expression data for all samples and leave it to data users to choose if and when to exclude these samples.

Next, we evaluated each dataset using the IQRray^33^ method, which identifies poor quality arrays by ranking all probe intensities in an array, computing the average rank for each probe set and returning a quality score for each sample in the dataset. Based on platform-specific quality thresholds from the authors’ prior analysis, we excluded samples that did not meet these thresholds. The IQRray method is specific to samples processed on Affymetrix platforms.

In steps four and five, we processed the metadata and expression data, respectively, for non-Affymetrix datasets. We created a separate script for each dataset because they were heterogeneous. For instance, the files were downloaded from different sources, and the variables we excluded differed widely in each of these datasets. We also had to separate the TCGA dataset into tumor-only and normal-only datasets. Otherwise, the tasks performed in these steps paralleled what we did in steps one and two.

In step six, we standardized the names of individual samples, ensuring consistency with capitalization and removing suffixes. We also removed the control probeset values from the Affymetrix datasets. Next we filtered out poor quality arrays identified by the IQRray method. To ensure consistency, we matched the samples in the expression data to samples in the metadata; thus the samples that were filtered by the IQRray method were also removed from the metadata. We also excluded any gene-expression samples for which metadata were missing. Finally, because some datasets used Entrez Gene identifiers while others used HUGO Gene Nomenclature Committee (HGNC)^34^ identifiers, we used the biomaRt^35,36^ package from Bioconductor^37^ to map gene identifiers across Entrez Gene, HGNC, and Ensembl^38^. When a single gene symbol was associated with multiple Ensembl identifiers, we retained genes from the main chromosomes and excluded gene identifiers from haplotypic regions. While this removed most duplicated genes, some remained. Through manual inspection, we found that the first identifier (sorted alphanumerically) was the most appropriate identifier, so we retained those. We determined that these identifiers were most appropriate by looking up each gene in The Human Gene Database^39^. This database has a section for each gene named “External Ids” where different identifiers are listed. We found that the first identifiers matched those listed in this section. We also excluded data for which any gene identifier was missing.

### Thesaurus mapping

We summarized the metadata variables into a single spreadsheet. For each variable, we used the variable name and data-value summaries to guide our interpretations of the semantic meaning. We manually mapped the metadata variables to NCIT standardized terms, using version:22.11d (release date: 2022-11-28). This was accessed via the NCI Term browser^40^. NCIT terms typically have synonyms and parent-child relationships, which proved useful in identifying the most appropriate term(s) for each variable. For this process, there were two curators (authors ION and SRP). The curators worked independently to match each variable name and value to respective NCIT terms. At the end of this process, the curators discussed the mappings jointly and addressed differences in interpretation. When we were uncertain about the semantic meaning, we searched the journal article associated with the dataset (if there was one) to see if any relevant explanations were available to decipher the meaning. These searches usually focused on the Methods sections or supplementary materials. For some datasets, we found relevant information in the supplementary files section on GEO. For variables that remained unclear, a practicing medical oncologist (author GC) provided suggestions. We excluded any variables that still remained unclear.

The spreadsheet has two columns: “NCIT_Term_field_name” and “NCIT_Term_values,” which contain the NCIT preferred name(s) and unique identifier(s) that we selected for each metadata variable name and its associated values, respectively. The former typically reflects the biomedical context of the variable; the latter typically indicates the manner in which the values were represented (e.g., units of measurement). In many instances, a single NCIT term was insufficient to convey the semantic meaning, so we used multiple NCIT terms. For example, for the variable *pr_status_by_ihc*, we used NCIT terms “Progesterone Receptor Status” and “Immunohistochemical Test Result”. In most cases, we retained the original metadata variable names; however, in a few cases, the names were particularly generic (e.g., *title* or *description*), so we renamed them. In other cases, a single column had multiple values per cell, so we split the column and named the new columns appropriately.

## Availability of data and materials

All gene expression data, metadata, and computer scripts generated during the current study are in a publicly accessible Open Science Framework (OSF)^41^ repository (accessible from https://osf.io/eky3p). The list of datasets is summarized in Table 1.

**Table 1:** Gene-expression data sources. This table summarizes all gene-expression datasets that we found in the public domain related to breast cancer.

| Author Name | Pubmed ID | Data Source | Number of samples* | Platform |
| --- | --- | --- | --- | --- |
| Zhou et al.<sup>47,48</sup> | 17407600, 18631401 | GSE7378 | 52 (54) | Affymetrix GeneChip HT-HG_U133A Early Access Array |
| Yau et al.<sup>48,49</sup> | 17850661, 18631401 | GSE8193 | 45 (47) | Affymetrix GeneChip HT-HG_U133A Early Access Array |
| Chin et al.<sup>50</sup> | 17157792 | E-TABM-158 | 129 (130) | Affymetrix GeneChip HT-HG_U133A Early Access Array |
| Knudsen et al.<sup>51</sup> | 22134623 | GSE33692 | 45 | Affymetrix Human Exon 1.0 ST Array [transcript (gene) version] |
| Tofigh et al.<sup>52</sup> | 25284793 | GSE58644 | 321 | Affymetrix Human Gene 1.0 ST Array [transcript (gene) version] |
| Winslow et al. | No associated publication found | GSE59772 | 9 | Affymetrix Human Gene 1.0 ST Array [transcript (gene) version] |
| Lehmann et al.<sup>53</sup> | 27310713 | GSE81838 | 20 | Affymetrix Human Gene 1.0 ST Array [transcript (gene) version] |
| Rebollar-Vega et al. | No associated publication found | GSE86374 | 159 | Affymetrix Human Gene 1.0 ST Array [transcript (gene) version] |
| Gregory et al.<sup>54</sup> | 31248446 | GSE118432 | 41 (42) | Affymetrix Human Gene 1.0 ST Array [transcript (gene) version] |
| Chang et al.<sup>55</sup> | 15718313 | GSE6434 | 19 (24) | Affymetrix Human Genome U95 Version 2 Array |
| Lee et al.<sup>56</sup> | 22751464 | GSE41194 | 49 (50) | Affymetrix Human Genome U95 Version 2 Array |
| Lee et al.<sup>56</sup> | 22751464 | GSE41196 | 11 (14) | Affymetrix Human Genome U95 Version 2 Array |
| Lee et al.<sup>56</sup> | 22751464 | GSE41197 | 16 (18) | Affymetrix Human Genome U95 Version 2 Array |
| Farmer et al.<sup>57</sup> | 15897907 | GSE1561 | 49 | Affymetrix Human Genome U133A Array |
| Wang et al.<sup>58</sup> | 15721472 | GSE2034 | 286 | Affymetrix Human Genome U133A Array |
| Minn et al.<sup>59,60</sup> | 16049480,<br>23982961 | GSE2603 | 121 | Affymetrix Human Genome U133A Array |
| Sotiriou et al.<sup>61</sup> | 16478745 | GSE2990 | 189 | Affymetrix Human Genome U133A Array |
| Karn et al.<sup>62–67</sup> | 17317819,<br>18058224,<br>19272155,<br>19455418,<br>21203899,<br>25274406 | GSE4611 | 216 (218) | Affymetrix Human Genome U133A Array |
| Minn et al.<sup>68</sup> | 17420468 | GSE5327 | 28 (58) | Affymetrix Human Genome U133A Array |
| Boersma et al.<sup>69–71</sup> | 17999412,<br>19225562,<br>20978357 | GSE5847 | 95 | Affymetrix Human Genome U133A Array |
| Desmedt et al.<sup>72,73</sup> | 17545524,<br>25788628 | GSE7390 | 198 | Affymetrix Human Genome U133A Array |
| Tripathi et al.<sup>74</sup> | 18058819 | GSE9574 | 29 | Affymetrix Human Genome U133A Array |
| Schmidt et al.<sup>75–78</sup> | 18593943,<br>20500820,<br>25485508,<br>33003293 | GSE11121 | 200 | Affymetrix Human Genome U133A Array |
| Zhang et al.<sup>79</sup> | 18821012 | GSE12093 | 134 (136) | Affymetrix Human Genome U133A Array |
| Zhang et al.<sup>80,81</sup> | 19573813,<br>25670079 | GSE14018 | 36 | Affymetrix Human Genome U133A Array |
| Emery et al.<sup>82</sup> | 19700746 | GSE16873 | 40 | Affymetrix Human Genome U133A Array |
| Symanns et al.<sup>83</sup> | 20697068 | GSE17705 | 296 (298) | Affymetrix Human Genome U133A Array |
| Miller et al.<sup>84,85</sup> | 20646288,<br>20697427 | GSE20181 | 147 (176) | Affymetrix Human Genome U133A Array |
| Shi et al.<sup>86,87</sup> | 20064235,<br>20676074 | GSE20194 | 251 (278) | Affymetrix Human Genome U133A Array |
| Tabchy et al.<sup>88,89</sup> | 20829329,<br>23185353 | GSE20271 | 156 (178) | Affymetrix Human Genome U133A Array |
| Graham et al.<sup>90</sup> | 20197764 | GSE20437 | 42 | Affymetrix Human Genome U133A Array |
| Graham et al.<sup>91</sup> | 21059815 | GSE21947 | 30 | Affymetrix Human Genome U133A Array |
| Iwamoto et al.<sup>89,92</sup> | 21191116, 23185353 | GSE22093 | 97 (103) | Affymetrix Human Genome U133A Array |
| Iwamoto et al.<sup>92</sup> | 21191116 | GSE23988 | 56 (61) | Affymetrix Human Genome U133A Array |
| Creighton et al.<sup>93</sup> | 21750966 | GSE24185 | 91 (103) | Affymetrix Human Genome U133A Array |
| Hatzis et al.<sup>94,95</sup> | 21558518, 36293478 | GSE25055 | 295 (310) | Affymetrix Human Genome U133A Array |
| Hatzis et al.<sup>94-96</sup> | 21558518, 24337596, 36293478 | GSE25065 | 184 (198) | Affymetrix Human Genome U133A Array |
| Karn et al.<sup>97-100</sup> | 21978456, 22220191, 21741824, 26484129, | GSE31519 | 67 | Affymetrix Human Genome U133A Array |
| Bianchini et al.<sup>101,102</sup> | 20805453, 22235097 | GSE32518 | 73 (74) | Affymetrix Human Genome U133A Array |
| Nagalla et al.<sup>103</sup> | 23618380 | GSE45255 | 135 (139) | Affymetrix Human Genome U133A Array |
| Rody et al.<sup>67,104-107</sup> | 18340528, 19054665, 23135572, 24737166, 25274406, | GSE46184 | 74 | Affymetrix Human Genome U133A Array |
| Casey et al.<sup>108</sup> | 18373191 | GSE10797 | 25 (66) | Affymetrix Human Genome U133A 2.0 Array |
| Reyngold et al.<sup>109,110</sup> | 25083786, 28813674 | GSE57968 | 71 (72) | Affymetrix Human Genome U133A 2.0 Array |
| Lu et al.<sup>111</sup> | 18297396 | GSE5460 | 127 (129) | Affymetrix Human Genome U133 Plus 2.0 Array |
| Turashvili et al.<sup>112</sup> | 17389037 | GSE5764 | 30 | Affymetrix Human Genome U133 Plus 2.0 Array |
| Richardson et al.<sup>113</sup> | 16473279 | GSE7904 (an extension of GSE3744) | 62 | Affymetrix Human Genome U133 Plus 2.0 Array |
| Karnoub et al.<sup>114</sup> | 17914389 | GSE8977 | 22 | Affymetrix Human Genome U133 Plus 2.0 Array |
| Loi et al.<sup>115,116</sup> | 18498629,<br>20479250 | GSE9195 | 77 | Affymetrix Human Genome U133 Plus 2.0 Array |
| Budczies et al.<sup>117</sup> | 21339180 | GSE11001 | 14 (30) | Affymetrix Human Genome U133 Plus 2.0 Array |
| Creighton et al.<sup>118</sup> | 19666588 | GSE10281 | 36 | Affymetrix Human Genome U133 Plus 2.0 Array |
| Chen et al.<sup>119</sup> | 19266279 | GSE10780 | 179 (185) | Affymetrix Human Genome U133 Plus 2.0 Array |
| Pedraza et al.<sup>120</sup> | 20029976 | GSE10810 | 58 | Affymetrix Human Genome U133 Plus 2.0 Array |
| Bos et al.<sup>121</sup> | 19421193 | GSE12276 | 204 | Affymetrix Human Genome U133 Plus 2.0 Array |
| Hoeflich et al.<sup>122,123</sup> | 19567590,<br>21673316 | GSE12763 | 30 | Affymetrix Human Genome U133 Plus 2.0 Array |
| Marty et al.<sup>124 125</sup> | 19055754,<br>21047405 | GSE13787 | 23 | Affymetrix Human Genome U133 Plus 2.0 Array |
| Zhang et al.<sup>80,81</sup> | 19573813,<br>25670079 | GSE14017 | 29 | Affymetrix Human Genome U133 Plus 2.0 Array |
| Desmedt et al.<sup>126</sup> | 19573224 | GSE16391 | 55 | Affymetrix Human Genome U133 Plus 2.0 Array |
| Li et al.<sup>127–130</sup> | 20098429,<br>20189874,<br>21422418,<br>26484051 | GSE16446 | 120 | Affymetrix Human Genome U133 Plus 2.0 Array |
| Sircoulomb et al.<sup>131</sup> | 20932292 | GSE17907 | 55 (109) | Affymetrix Human Genome U133 Plus 2.0 Array |
| Korde et al.<sup>132</sup> | 20012355 | GSE18728 | 61 | Affymetrix Human Genome U133 Plus 2.0 Array |
| Li et al.<sup>127,128,133</sup> | 20098429,<br>20100965,<br>20189874 | GSE18864 | 71 | Affymetrix Human Genome U133 Plus 2.0 Array |
| Li et al.<sup>127</sup> | 20098429 | GSE19615 | 115 | Affymetrix Human Genome U133 Plus 2.0 Array |
| Lin et al.<sup>134</sup> | 19967557 | GSE19697 | 24 | Affymetrix Human Genome U133 Plus 2.0 Array |
| Bauer et al.<sup>135</sup> | 20062080 | GSE20086 | 12 | Affymetrix Human Genome U133 Plus 2.0 Array |
| Kao et al.<sup>136</sup> | 21501481 | GSE20685 | 327 | Affymetrix Human Genome U133 Plus 2.0 Array |
| Dedeurwaerder et al.<sup>137</sup> | 21910250 | GSE20711 | 90 | Affymetrix Human Genome U133 Plus 2.0 Array |
| Kretschmer et al.<sup>138,139</sup> | 21314937, 21468687 | GSE21422 | 19 | Affymetrix Human Genome U133 Plus 2.0 Array |
| Sabatier et al.<sup>140,141</sup> | 20490655, 22110708 | GSE21653 | 266 | Affymetrix Human Genome U133 Plus 2.0 Array |
| Bauer et al.<sup>142–144</sup> | 20068102, 20878462, 21633166 | GSE22513 | 28 | Affymetrix Human Genome U133 Plus 2.0 Array |
| Hawthorn et al.<sup>145</sup> | 20799942 | GSE22544 | 19 (20) | Affymetrix Human Genome U133 Plus 2.0 Array |
| Bekhouché et al.<sup>141,146,147</sup> | 21339811, 22110708, 22666420 | GSE23720 | 197 (370) | Affymetrix Human Genome U133 Plus 2.0 Array |
| Latimer et al.<sup>148</sup> | 21118987 | GSE25407 | 10 | Affymetrix Human Genome U133 Plus 2.0 Array |
| Planche et al.<sup>149</sup> | 21611158 | GSE26910 | 12 (24) | Affymetrix Human Genome U133 Plus 2.0 Array |
| LaBreche et al.<sup>150</sup> | 21781289 | GSE27562 | 161 (162) | Affymetrix Human Genome U133 Plus 2.0 Array |
| Lehmann et al.<sup>144</sup> | 21633166 | GSE28821 | 89 | Affymetrix Human Genome U133 Plus 2.0 Array |
| Cuadros et al.<sup>151,152</sup> | 22622817, 22832278 | GSE29431 | 60 (66) | Affymetrix Human Genome U133 Plus 2.0 Array |
| Jones et al. | No associated publication found | GSE31138 | 6 | Affymetrix Human Genome U133 Plus 2.0 Array |
| Harvel et al.<sup>153</sup> | 23479404 | GSE31192 | 33 | Affymetrix Human Genome U133 Plus 2.0 Array |
| Sabatier et al.<sup>141</sup> | 22110708 | GSE31448 | 357 | Affymetrix Human Genome U133 Plus 2.0 Array |
| Miyake et al.<sup>154</sup> | 22320227 | GSE32646 | 115 | Affymetrix Human Genome U133 Plus 2.0 Array |
| Clarke et al.<sup>155</sup> | 23740839 | GSE42568 | 115 (121) | Affymetrix Human Genome U133 Plus 2.0 Array |
| Metzger-Filho et al.<sup>156</sup> | 23990869 | GSE43365 | 111 | Affymetrix Human Genome U133 Plus 2.0 Array |
| D'Alfonso et al.<sup>157</sup> | 23774991 | GSE47109 | 229 (246) | Affymetrix Human Genome U133 Plus 2.0 Array |
| Huang et al.<sup>158</sup> | 24098497 | GSE48390 | 80 (81) | Affymetrix Human Genome U133 Plus 2.0 Array |
| Prat et al.<sup>159</sup> | 24443618 | GSE50948 | 78 (156) | Affymetrix Human Genome U133 Plus 2.0 Array |
| Fumagalli et al. | No associated publication found | GSE58984 | 94 | Affymetrix Human Genome U133 Plus 2.0 Array |
| Aswad et al.<sup>160,161</sup> | 26474389, 26517092 | GSE61304 | 62 | Affymetrix Human Genome U133 Plus 2.0 Array |
| Burstein MD et al.<sup>162,163</sup> | 25208879, 26921331 | GSE76275 | 265 | Affymetrix Human Genome U133 Plus 2.0 Array |
| Brouwers et al.<sup>164</sup> | 28673354 | GSE90521 | 17 | Affymetrix Human Genome U133 Plus 2.0 Array |
| Santucci-Pereira et al.<sup>165</sup> | 30922380 | GSE111662 | 27 | Affymetrix Human Genome U133 Plus 2.0 Array |
| Dhage et al.<sup>166</sup> | 30325954 | GSE120129 | 107 (108) | Affymetrix Human Genome U133 Plus 2.0 Array |
| Hartung et al.<sup>167</sup> | 35082572 | GSE167213 | 87 (124) | Affymetrix Human Genome U133 Plus 2.0 Array |
| Lehmann et al.<sup>144</sup> | 21633166 | GSE28796 | (24) | Affymetrix Human Genome U133 Plus 2.0 Array |
| Jose et al.<sup>168</sup> | 30169535 | GSE93332 | (8) | Affymetrix Human Genome U133 Plus 2.0 Array |
| Pawitan et al.<sup>169,170</sup> | 16280042, 16813654 | GSE1456 | 317 (318) | Affymetrix Human Genome U133A / U133B Array |
| Miller et al.<sup>171–173</sup> | 16141321, 29136509, 28972043 | GSE3494 | 501 (502) | Affymetrix Human Genome U133A / U133B Array |
| Ivshina et al.<sup>174</sup> | 17079448 | GSE4922 | 570 (578) | Affymetrix Human Genome U133A / U133B Array |
| Loi et al.<sup>115,116,175,176</sup> | 17401012, 18498629, 20479250, 19552798 | GSE6532 | 741 | Affymetrix Human Genome U133A / U133B / U133 Plus 2.0 Array |
| Rahman et al.<sup>25,26</sup> | 26209429, 23000897 | GSE62944 (TCGA) | 1171 | Illumina Genome Analyzer |
| Brueffer et al.<sup>177–179</sup> | 32913985, 32926574, 33937624 | GSE81538 | 405 | Illumina HiSeq 2000 |
| Brueffer et al.<sup>177–180</sup> | 32913985, 32926574, 33937624, 35304506 | GSE96058 | 3273 (3409) | Illumina HiSeq 2000, Illumina NextSeq 500 |
|  | No associated publication | https://dcc.icgc.org/releases/current/Projects/BRCA-KR (ICGC) | 50 | Illumina HiSeq |
| Curtis et al.<sup>24</sup> | 22522925 | https://www.cbioportal.org/study/summary?id=brca_metabric (METABRIC) |  | Illumina Human HT-12 v3 Expression Beadchips |

\* When the sample size was different after filtering, we indicate the original sample size in parenthesis.

## Discussion

Huge sums are budgeted each year by the National Institutes of Health and other grant-awarding agencies to generate gene-expression data and accompanying metadata. Additionally, researchers put much time and effort into generating the original data. Placing the data in the public domain is invaluable for research, but that alone is not enough. To move the field forward, we need compendia of datasets that follow common themes. By reprocessing, curating, and sharing these breast-cancer datasets, we are able to save time that other researchers would have used in completing the same steps, thus reducing duplicated efforts. These efforts make the data more accessible to researchers who lack the expertise to complete these reprocessing and curation steps. Accordingly, we have sought to increase the benefit to the public of prior investments.

A major takeaway from this project is the number of samples that we have compiled. This resource includes 102 datasets, representing 17,151 patient samples. While groups like TCGA^25^ have generated large data collections (11,000 tumor samples across 33 cancer types), we have collated data from multiple sources and have focused solely on breast cancer. Projects like *recount3*^42^ focus on RNA sequencing data for a wide range of biomedical conditions. Out of necessity, that project provides relatively little metadata, which is uncurated. We have taken considerable time to clean and manually curate the metadata variables in the datasets we reprocessed because it is difficult for many researchers to decipher the semantic meaning of a variable solely by its name or contents. Human review is essential (and time consuming). We envision that researchers will search our compendium for datasets associated with given NCIT term(s) and perform diverse types of meta-analyses using those data. Supplementary File 1 contains these NCIT term mappings.

To ensure reproducibility, we tested our scripts using an iterative process. The scripts have been configured to be executed within Docker containers so that they can be re-executed more easily by others on different operating systems and obtain the same outputs. Finally, our output files are simple text files and thus are programming-language agnostic. Other research groups have created valuable packages like the MetaGxBreast^19^ and curatedBreastData^20^ packages. These packages are accessible using the R programming language and are implemented within the Bioconductor framework, which is used heavily in the biomedical community^43^, but the outputs from our scripts can be accessed on any platform.

Although our compendium consists of many datasets relevant to a common biomedical theme, care must be taken when combining datasets. Systematic differences between data-generation platforms, laboratory equipment, and environmental conditions can bias analytical results^44^. We did not adjust for these differences but rather leave it to consumers of the data to identify scenarios and techniques for adjusting the data. Analyses that combine *inferences* across datasets—rather than combining data directly—may be less prone to such biases.

We mapped the metadata to NCIT terms but did not standardize the field names or values in the text files. Therefore, when combining datasets, researchers may wish to perform this step. For instance, one dataset might have a column that indicates estrogen-receptor status using values of "1" or "0", whereas another dataset might represent the same information using values of "positive" and "negative" or "P" and "N". To provide insight into the frequency with which biomedical concepts are represented in the metadata, we summarized the NCIT terms mapped to the metadata fields. Summaries for the NCIT terms used to describe field names can be found in Supplementary Files 2 and 3. Some of the most commonly observed terms for field names were Estrogen Receptor Status (n = 73), Event-Free Survival (n = 63), Disease Stage Qualifier (n = 52), Progesterone Receptor Status (n = 46) and HER2/Neu Status (n = 43). Thus, even though some variables are quite common, these results illustrate that breast-cancer studies differ considerably in the metadata collected. Some terms like sex (n = 15) and race (n = 11) were less common than we expected. Even though breast cancer largely affects females, approximately 1% of cases affect males^45^; recording research participants’ sex would make it possible to learn more about differences between male and female cases. Collecting race information may make it possible to shed more light on mechanisms that drive differential responses to treatments across racial groups^46^. Summaries for the NCIT terms used to describe data values can be found in Supplementary Files 4 and 5. Across all metadata fields, the most commonly observed value was “binary” (n = 138). Researchers used either 0 or 1 to represent many variables, including estrogen-receptor status, tumor size and lymph node status. In other cases, terms like Estrogen Receptor Positive and Estrogen Receptor Negative were used. In these cases the values can be more easily interpreted, whereas binary values must be interpreted in combination with field names. In future studies it may be beneficial to use more explicit terms to describe these values.

The large number of unique field names and values indicates the huge diversity in data that researchers collect. However, only 23% of terms (25% for values) occurred in 5 or more datasets. With this low concordance across datasets, it may be difficult to draw broad conclusions across the datasets. In future studies, it would be beneficial to collect a common set of variables across studies (although researchers might collect additional variables that are specific to each study).

In addition to aiding breast-cancer researchers in identifying datasets and samples associated with research topics of interest, biomedical-semantics researchers can use the ontology mappings for developing and testing algorithms that automate the ontology-mapping process. Although we were careful to ensure consistency when annotating the metadata variables, this process is subjective in nature. In many cases, several different NCIT terms could have been used to represent a given biomedical concept. In nearly every case, more generic (parent) or specific (child) terms could have been used as alternatives. The NCI Term Browser provides mappings between terms and links to other ontologies, thus making it possible for researchers to use our annotations as a starting point to find alternative terms.

## Conclusions

This standardized resource has the potential to reduce time that breast-cancer researchers spend on finding and preparing publicly available gene-expression data and allow them to focus more on answering biomedical questions. We have previously^9^ surveyed the literature on types of analyses that have been commonly performed using gene-expression data from breast-cancer patients. Examples include predicting patient outcomes, defining breast-cancer subtypes, and identifying differentially expressed genes. We believe this resource will help in making these types of studies easier and to have more statistical power.

## Supplementary Data

Supplementary File 1 is a summary of the metadata in which the variables have been mapped to the NCIT terms.

Supplementary FIle 2 is a frequency table of the individual NCIT terms mapped to field names.

Supplementary FIles 3 is a frequency table of combinations of NCIT terms mapped to field names.

Supplementary FIles 4 is a frequency table of the individual NCIT terms mapped to data values.

Supplementary FIles 5 is a frequency table of combinations of NCIT terms mapped to data values.

## Notes

https://osf.io/eky3p

### Competing Interest Statement

The authors have declared no competing interest.

### Summary of Updates

We fixed some formatting problems with the supplementary files.

